# Potential anti-adipogenic activity of *Calligonum comosum* cuminaldehyde on mouse 3T3-preadipocytes

**DOI:** 10.1101/2020.07.02.184200

**Authors:** Mohammad G Mohammad, Ahmed El-Serafi, Mohamed I. Madkour, Abeer Alhabshi, Ansar Wadea, Rola Abu Jabal, P Divyasree, Sameh S.M. Soliman

## Abstract

Obesity is a medical condition associated with serious medical and psycho-social consequences and an augmented body fat mass. Several compounds were suggested to counteract obesity and fat accumulation with variable degrees of success. Searching for a safe and effective anti-adipogenic substance, we found that cuminaldehyde-rich essential oil extracted from *Calligonum comosum* potentially mediate activities. The results showed that *C. comosum* essential oil and its major component cuminaldehyde, selectively caused significant reduction in the viability of 3T3-L1 cells when compared to fibroblasts. Furthermore, cuminaldehyde caused significant reduction in the lipid content, glucose uptake and levels of both triglycerides and cholesterol in adipocytes. Moreover, the formation of 3D-adipocyte pellets in the presence of cuminaldeyde was affected. Adipocytes matured in the presence of cuminaldehyde have significant reduction in the expression of adipocyte-specific transcripts, CAAT-enhancer binding protein alpha (CEBPa) and Peroxisome proliferator-activated receptor gamma (PPARg). Taken together, these results demonstrate a potential inhibitory role of cuminaldehyde extracted from *C. comosum* oil on lipid accumulation. Consequently, cuminaldeyde can be considered as a new potential anti-adipogenic agent for prevention and treatment of obesity.

## Introduction

Obesity is a medical condition in which the body fat mass is increased to a level that can adversely affect human health. According to the World Health Organization (WHO), obesity has tripled since 1975 and 39% of adults aged 18 years and above were overweight in 2020 and 13% were obese [1]. Obesity is considered as a risk factor for life-threatening conditions, including cardiovascular diseases [2], musculoskeletal disorders [3] and some types of cancer [4]. A fundamental cause of obesity is an energy imbalance between calories consumption and exhaustion [5]. Obesity is largely preventable and can be managed by applying several strategies including the use of therapeutic compounds [6]. A key strategy in reducing obesity is dampening adipocyte differentiation, reducing adipocyte hypertrophy and hyperplasia, as well as antagonizing fat accumulation [7]. Several therapeutic compounds were developed in order to control general body obesity, as well as localized fat accumulation [8]. On the other hand, local communities accumulated considerable experience for the use of native growing plants as a treatment ofadiposity. For example, compounds such as scoparone isolated from Chinese herb *Artemisia scoparia* [9], quinoa saponins [10], myricetin [11], 4-hydroxyderricin and xanthoangelol from Japanese Ashitaba (*Angelica keiskei*) [12] have shown significant interference with adipocyte functions. *Calligonum comosum L*. has been employed by local communities in the United Arab Emirates (UAE) for decades in their foods for the treatment of several gastric and skin illnesses [13,14]. Various pharmacological activities of the plant were also reported, including anti-ulcer, anti-inflammatory and anti-cancer activities [15,16]. *C. comosum* is known as a source of volatile essential oil [15,17]. The purpose of this study was to investigate the anti-adipogenic activity of *C. comosum* oil and to determine a new lead compound that might interfere with adipocytes differentiation and metabolism.

## Experimental procedures

### Plant materials

*C. comosum* L. (*Polygonaceae*) plant materials (stems and roots) were collected during December 2016 from the desert of Sharjah (Coordinates 25.3284° N, 55.5123° E) [18], United Arab Emirates. The plants were collected and identified according to the international standard of the plant taxonomy and by Prof. Ali El-Keblawy (Department of Biology, College of Sciences, Sharjah) and voucher specimen were deposited at University of Sharjah herbarium. There is no specific permission required for collecting plant samples from the open desert of UAE for scientific purposes. The collected plant samples are not classified as endangered species and not under protection protocol.

### Plant essential oil preparation and analysis

Air-dried plant stems were grinded, and the essential oil was extracted by steam distillation method according to [19,20]. Briefly, 200gm plant stem powder was extracted with water steam at 100°C for 3 hr. The condensed oil was collected and dried over anhydrous sodium sulphate to yield 250 µL. The oil was diluted in chloroform in a ratio of 1:100 prior to GC-MS analysis. The composition of essential oil was analyzed and identified according to Soliman *et al*., 2017 [21] using the Agilent Gas Chromatography-Mass Spectrometry (GC-MS). The identified compounds including the target compound, cuminaldehyde, were described in Soliman *et al*., 2017 [19] in both tabular and chromatogram forms.

### Fractionation-based bioassay and partial purification of cuminaldehyde

Identification of the major anti-adipogenic compound in the plant essential oil extract was performed according to Soliman *et. al*, [20]. Briefly, the plant essential oil was separated on analytical TLC plates (Sigma-Aldrich, Germany) and developed by petroleum ether: acetone (5:1) as solvent system. The plate was visualized under UV lamb using short wavelength (250nm). The major bright bands were scrapped off the plate and each fraction was tested for its anti-adipogenic activity using 3T3 cells. The fraction (# 3) containing the active compound, cuminaldehyde, was confirmed by GC-MS. The percentage of cuminaldehyde in the fraction was > 50%. The fractionation TLC and corresponding GC-MS chromatogram was described in Soliman *et al*., 2020 [20].

### Cell culture

The 3T3-L1 mouse pre-adipocyte cell line was originally developed by clonal expansion from murine Swiss 3T3 cells. Normal fibroblast cells line (F180) was employed as control. Both cell lines were cultured in the Dulbecco’s Modified Eagle Media (DMEM; Sigma-Aldrich, Germany) supplemented with 25 mM glucose, 10% Fetal Bovine Serum (FBS; Sigma-Aldrich, Germany) and 1% penicillin-streptomycin (Sigma-Aldrich, Germany) and maintained at 37°C in a humidified atmosphere of 5% CO_2_.

### MTT cell viability assay and determination of LD_50_

The reduction of yellow tetrazolium salt 3-(4, 5-dimethylthiazol-2-yl)-2,5-diphenyltetrazolium bromide (MTT; Sigma-Aldrich, Germany) was used to measure cellular metabolic activity as a proxy for cell viability [19,22]. To measure the lethal dose that is required to induce 50% death of the cells (LD50), 3T3 and F10 cell lines were cultured till reached 80% confluency. A 96-well plate was seeded with 4000 cells per 100μl media and incubated at 37°C for 24 hrs. A serial dilution of *C. comosum* oil extract, its separated fractions, or cuminaldehyde-containing fraction were added onto the cells and incubated for further 24 hrs. Freshly prepared MTT solution (5 mg/ml) was added to each well (20 μl per well) and followed by incubation for 2 hrs at 37°C. The supernatant were then removed and 100 μl DMSO was added and incubated until formazan violet crystals were developed and the OD_540_ was measured.

### Pre-adipocyte cell differentiation

Differentiation of pre-adipocytic 3T3 cells was followed a published protocol by Vishwanath et.at., [23]. Briefly, 2×10^4^ pre-3T3 cells were seeded in a 12-well plate till reached confluency level. The medium was then supplemented with 10 µM dexamethasone [Sigma-Aldrich, Germany], 0.5 mM 3-isobutyl-1-methylxanthine (IBMX; Calbiochem, La Jolla, CA), 100 µM indomethacin (Sigma-Aldrich, St. Louis, MO), and insulin-transferrin-sodium selenite (ITS) supplement to provide 10 µg/ml of insulin (Sigma-Aldrich, Germany). After three days, the media was replaced with fresh media supplemented with ITS only. The last step was repeated once more cycle after three day. Cultures were maintained in a humidified incubator containing 5% (v/v) CO_2_ at 37°C. The accumulation of lipid droplets and adipocytes growth was monitored over the cultivation time.

### Cholesterol, triglyceride and glucose-intake measurements

Post-differentiated adipocytes were incubated for 24 hrs with *C. comosum* cuminaldehyde fraction in 12-well plate at its LD_50_. Wells received no oil were employed as a negative controls. Cell lysates and culture supernatants were then obtained and stored at -80°C prior to testing. Ezymatic colorimetric assays were used to measure cholesterol, triglycerides and glucose (using Diasys colorimetric kits, Diagnostic systems) in comparison with provided known standard (DIASYS) [24,25].

### Nile red stain and flow cytometry

To quantify the changes in intracellular lipid droplets accumulation in differentiated adipocytes following the treatment for 24 hrs, Nile red stain was employed. Nile red stock solution (Sigma-Aldrich, Germany) was prepared in acetone and diluted in PBS at concentration of 0.5 μg/mL. The cells were then fixed with 4% paraformaldehyde. Fixed cells were stained with Nile red working solution (diluted 100× in 50 mM Tris/Maleate) for 5 min at room temperature. Stained cells were further examined under Olympus BX51 fluorescence microscope (Olympus Corporation, Tokyo, Japan) at excitation/emission wavelengths of 552/636 nm. Differential interference contrast (DIC) images were taken to produces contrast by visually displaying the refractive index gradients of different areas of 3T3 cells. Other sets treated similarly were gently scrapped after staining and positively stained cells were quantified using BD FACS Area III flow cytometer.

### RNA extraction, reverse transcriptase and semi-quantitative rT-PCR

To evaluate the gene expression related to the adipocyte differentiation, RNA was extracted from treated and non-treated cells by the column-based RNeasy Mini Kit (Qiagen), according to manufacturer’s instructions. Extracted RNA was quantified and reverse transcribed to cDNA using TruScript™ kit (Norgen, Thorold, Canada). For RT-PCR, cDNA was added as 100 ng/reaction to SYBR green based GoTaq^®^ qPCR Master Mix (Promega, Wisconsin, USA). The primer sequence for CEBPa was F: 5′-TTACAACAGGCCAGGTTTCC-3′ and R: 5′-GGCTGGCGACATACAGATCA-3, PPARγ was F: 5′-TTTTCAAGGGTGCCAGTTTC-3′ and R: 5′-AATCCTTGGCCCTCTGAGAT-3′. Beta actin was employed as housekeeping gene for normalization. The primer sequence was F: 5′-GACAACGGCTCCGGCATGTGCAAAG-3′ and R: 5′-TTCACGGTTGGCCTTAGGGTTCAG-3′. The polymerase chain reaction (PCR) were carried out using Rotor Gene Q (Qiagen, Hilden, Germany) with initial incubation at 95 °C for 15 min, followed by 40 cycles of 95 °C for 15 s, 55 °C for 30 s and 70 °C for 30 s. Gene expression was calculated after normalization to β-actin following the 2^−ΔΔCT^ method.

### Pre-adipocyte pellet cultures

To evaluate the anti-adipogenic effects of cuminaldehyde on adipose tissue, 3T3 adipocytes were transferred to three dimensions as pellet culture, which provides a more physiological environment for cell to cell interaction and differentiation. The procedure of adipocyte pellet cultures preparation was performed according to [26,27]. Briefly, 1×10^6^ 3T3 cells/pellet were suspended in 1 ml media and centrifuged for 10 min at 400xg, followed by media change for three times per week. The pellets were then treated with cuminaldehyde at LD50 for 24 hours.

### Statistical analysis

Values were presented as means ± standard error of mean (SEM). Experiments were repeated with 3-6 replicas per each experiment. Data was analyzed by T test using Graphpad Prism software. Statistical significance was considered at *P-*value < 0.05.

## Results

### *C. comosum* essential oil showed potential and safe anti-adipogenic activity

The effect of *C. comosum* essential oil on the viability of pre-adipocyte mouse cell line (3T3-L1) and normal fibroblasts cell line (F180) were determined using MTT assay. The results showed that *C. comosum* oil caused selective inhibitory activities on 3T3 when compared to normal F180 cells. The oil at concentrations 0.4-0.8 μg/ mL caused significant reduction (∼50%, *P-*value < 0.05) in the viability of 3T3 cells when compared to F180 (**Figure 1**). The results obtained indicated that *C. comosum* essential oil can be employed as potential anti-adipogenic and 0.4-0.8 μg/ mL is the concentration of choice for selective inhibition of 3T3 cells.

**Figure 1.**
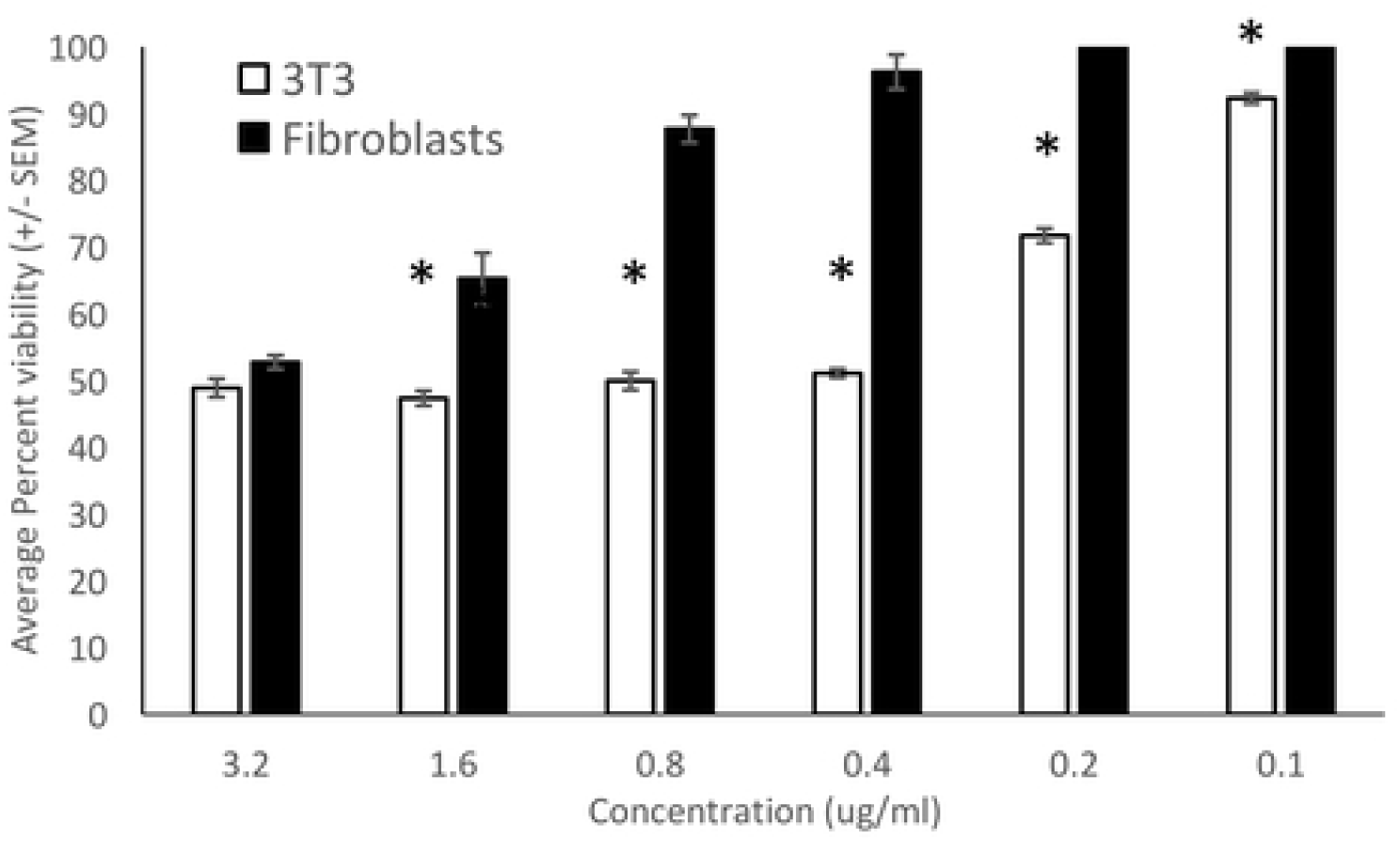
Effect on viability of cuminaldehyde rich oil separated from *C. comosum against* 3T3 and fibroblast. The viability of cuminaldehyde-treated and untreated cells was measured using the MTT assay. Shown is average of 3 separate experiments with 6 replicas per each experiment ± SEM. *: *p* value < 0.05.

### Cuminaldehyde was the oil major component responsible for the anti-adipogenic activity

The plant oil was subjected to fractionation-based 3T3 cell viability bioassay using MTT assay. The oil was fractionated on TLC into 5 fractions and each fraction was tested on 3T3 cells. Fraction number 3 that contained > 50% cuminaldehyde showed 90-100% killing activity when compared to other fractions, while its LD_50_ was determined as 0.4 μg/ mL (**Figure 1**).

Cuminaldehyde was further tested on the intracellular lipid accumulation of 8-day post-differentiated 3T3 adipocytes using Nile red staining. The results indicated that cuminaldehyde oil at the concentration of 0.4 μg/ mL caused significant reduction (25%, *p-*value =0.038) in the intracellular lipid content of 3T3 cells (**Figure 2A-D**). The concentration of both triglycerides and cholesterol from cell lysates and the level of glucose in media supernatants were measured. The results showed non-significant reduction in the triglyceride content (∼10% reduction, *P-* value =0.495), while showed 50% reduction in the cholesterol level contents at concentration 0.4 μg/ mL (*p-*value=0.009; **Figure 3A**). On the other hand, the glucose level in oil treatments was reduced by 40% (*p-*value=0.0012; **Figure 3B**).

**Figure 2.**
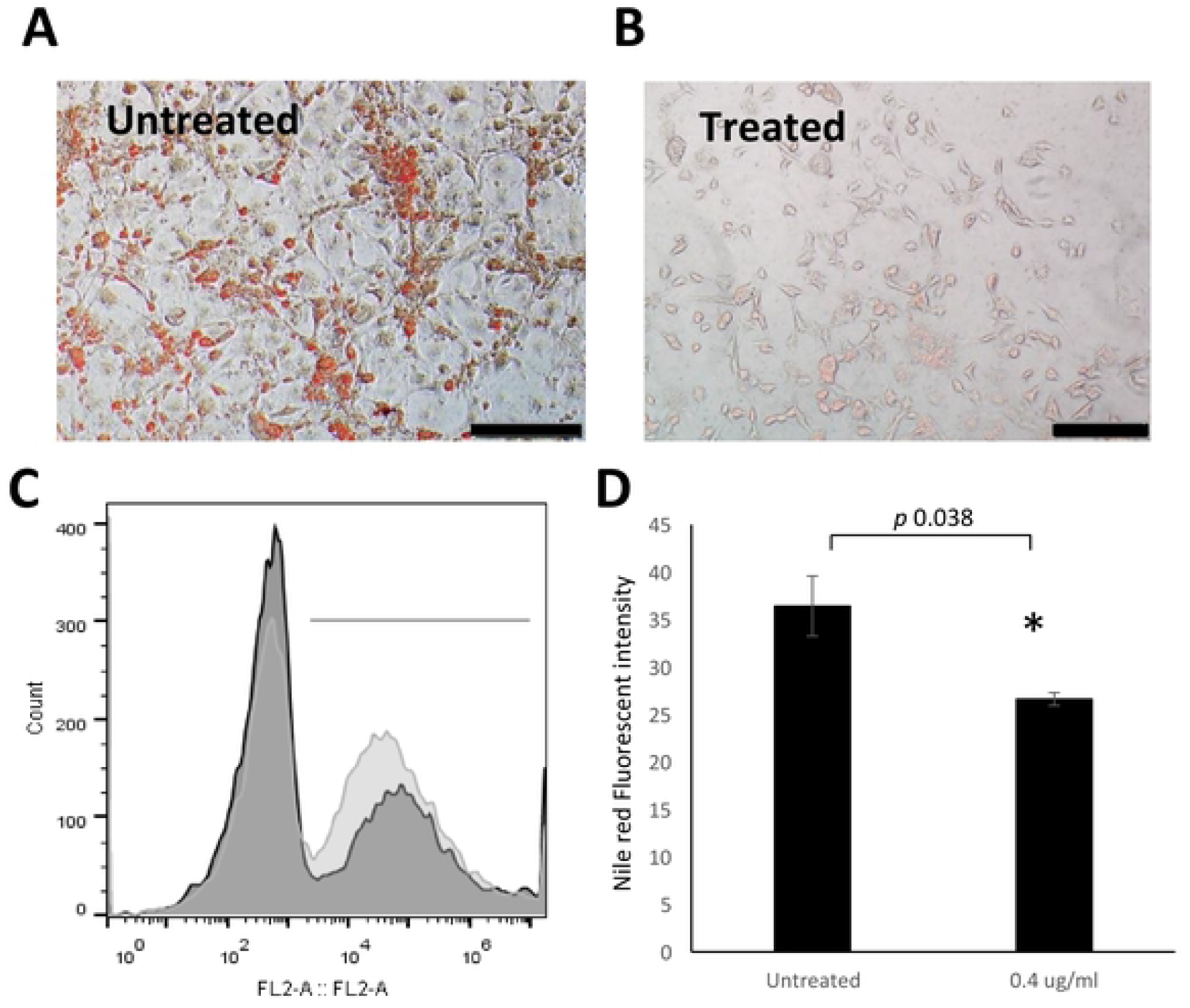
Reduction in the number of cells and the content of fat droplets within cuminaldehyde-treated 3T3 L1 cell line. Cuminaldehyde demonstrated significant reduction in the amount of fat droplet content within 3T3 L1 cell line after 24 hrs (**A** and **B**). 3T3 cells cultured in 6-well plates were stained with Nile red and the images taken by fluorescent microscope (FL-1 fluorescence and merged with DIC) where (**A**) represented control treatment and (**B**) represented treated cells with cuminaldehyde. (**C**) 3T3 cells were scraped, stained with Nile red, filtered and measured by BD FACS Area II flow cytometer. The histogram shows reduced fluorescence of treated cells (dark gray) compared to non-treated cells (light gray). (**D**) Nile red fluorescence intensity was significantly decreased in comparison to the untreated control. Fluorescence intensity averages ± SEM are shown. Scale bar=100μm, *: *p* value < 0.05.

**Figure 3.**
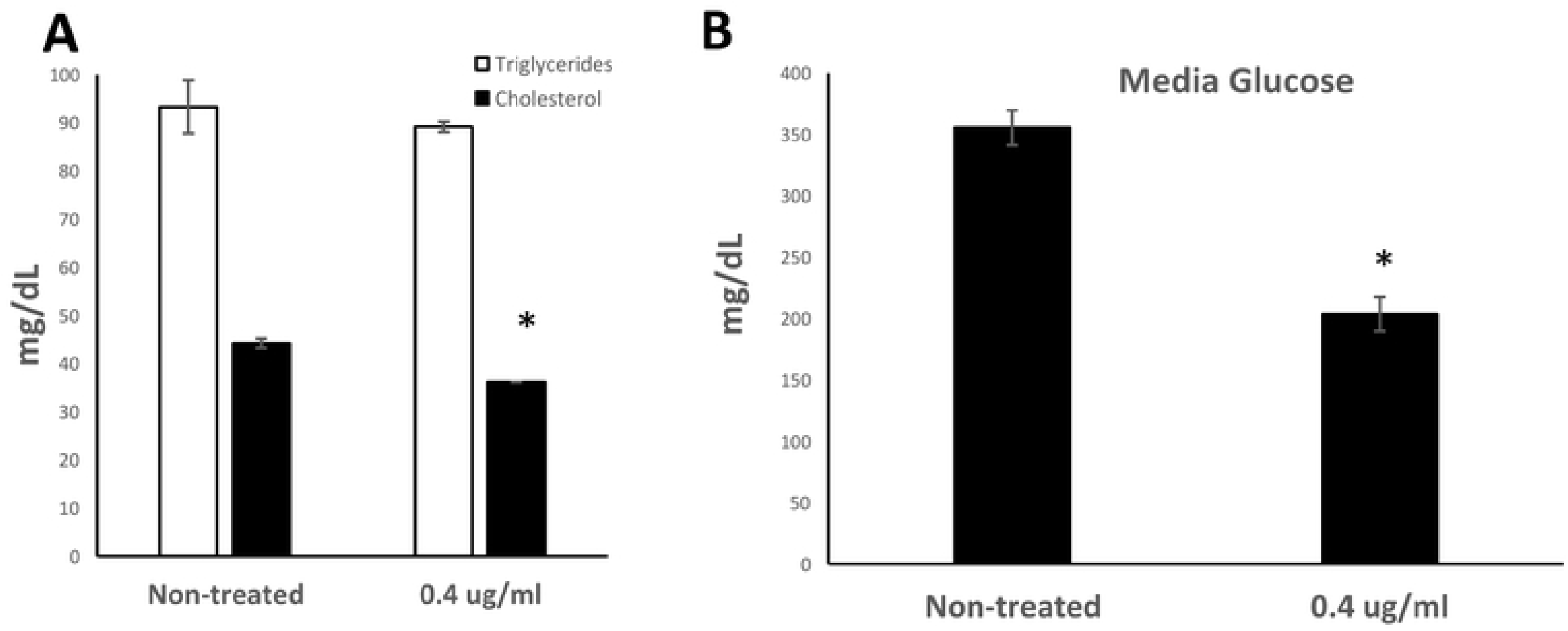
Effect of cuminaldehyde on the lipid and glucose metabolism of 3T3 cells. (**A**) Evaluation of triglycerides and cholesterol content in the cell lysate, is in response to treatment with cuminaldehyde showed decrease in the cholesterol content, in comparison to untreated control. (B) Similarly, the glucose content in the media was less in response to treatment. *: *p* value < 0.05.

Expression of the adipogenesis transcription factors CEBPa and PPARg is crucial for the adipocyte maturation [28]. Adding cuminaldehyde oil at concentration 0.4 μg/ mL caused significant reduction in the expression of CEBPα and PPARγ by approximately 50% and 70%, (*p-*value=0.01 and 0.0001, respectively; **Figure 4A**).

**Figure 4.**
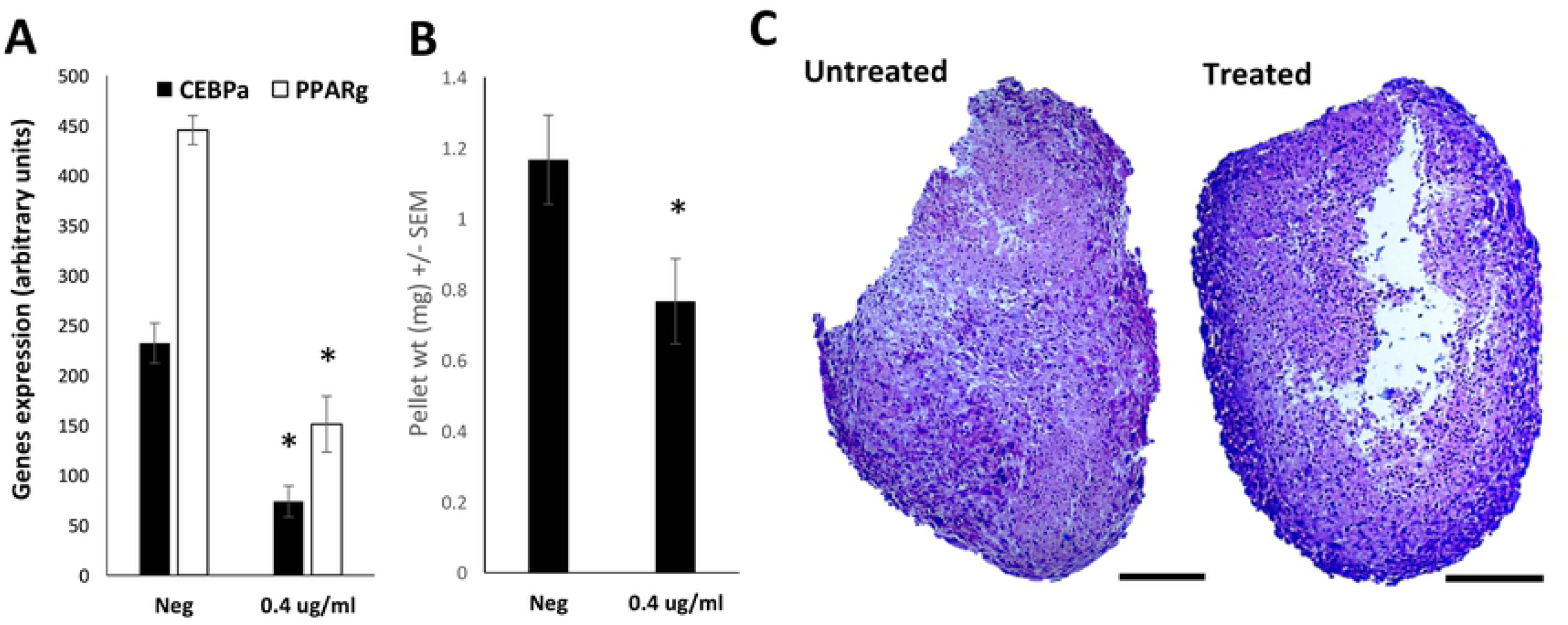
Cuminaldehyde significantly reduced the transcription of adipogenesis transcription factors and the weight of formed adipocyte pellets.. (**A**) The effects of cuminaldehyde on the expression of adipogenesis transcription factors CEBPa and PPARg. The number of mRNA transcripts of both genes is significantly reduced as a result of treatment. β-actin was employed for normalization. **(B)** Three-weeks old 3T3 cells were induced for the formation of 3D-culture pellets followed by treatment with cuminaldehyde compared to untreated control. Pellets weight measurements indicated a significant reduction in the weight of treated pellets. (**C**) Hematoxyline/ Eosin stained sections of treated and untreated pellets indicated interrupted core formation of the treated pellets. Scale bar= 200 um.. The results indicated as the mean ± SEM. *: *p* value < 0.05.

Additionally, cuminaldehyde caused reduction by 30% in the weight of the 3D adipogenic pellet (*P*-value=0.044; **Figure 4B**). Testing the histological structure of the formed adipocytes pellets indicated an absence of intact inner center as compared to the untreated controls (**Figure 4C**).

## Discussion

Adiposity is one of the world’s health problems that affects approximately third of the population all over the globe and can contribute to severity of many chronic diseases [29]. Several strategies and regimens have been established to control and reduce lipid accumulation, including those dependent on the inhibition of adipogenicity [30]. Strategies used to reduce adiposity can include controlling calories intake, induction of lipolysis, enhancement of adipocyte apoptosis and suppression of adipocyte differentiation [31]. Here in this study, we demonstrated the ability of pure cuminaldehyde and cuminaldehyde-rich oil isolated from, *C. comosum* as a natural product, to enhance adipocyte metabolism and hence induction of lipolysis.

General or local obesity can be a result of increase in the tissue fat mass which is primarily consists of mature differentiated adipocytes [32]. Thus, both lipolysis and inhibition of adipocyte differentiation can be considered as effective strategies in body weight loss or dissolution of local fat deposits. Consistently, our results indicated that cuminaldehyde caused cytotoxic effects on 3T3 cells. Inhibition of 3T3 is a known model associated with reduction in the tissues’ fat mass [33]. On the other hand, the cytotoxic effects of cuminaldehyde on 3T3 were accompanied by enhanced glucose intake and lipid turnover, which could be considered as an indicator of enhanced metabolism and/ or cells metabolic stress. Enhanced metabolism can maintain the body energy stores and hence regulate the storage and mobilization of fat stores [34]. In this regard, cuminaldehyde can be considered as a hypoglycemic inducing factor. The significant reduction in fat globules (stained with Nile red) due to the effect of cuminaldehyde, could suggest that the compound interfered with fat formation by holding the adipogenesis or reflecting the dissolution of fat in adipocytes. A dampened adipogenesis would result in reduced fat production and formation of intact adipose tissue. This observation was supported by the significant reduction in the expression of transcripts known to control adipocyte maturation, as a result of treatment with the compound [28]. These results can indicate that cuminaldehyde anti-adipogenic potential effects can be – at least partially-through antagonizing the expression of adipogenesis-related genes. The suppression effects of cuminaldehyde on adipocyte maturation are similar to inhibition effects of cocoa tea [35].

The process of adipogenesis is highly regulated by complex network including transcription factors, calories intake, fat used up and adipocyte maturation. *C. comosum* essential oil extract, and its major component cuminaldehyde, showed potential anti-adipogenic activities. Cuminaldehyde increased the metabolic activities of adipocytes and hence the consumption of stored lipids for energy use along with suppression of pre-adipocytes differentiation. The aforementioned double activities of cuminaldehyde by reducing lipid content can be employed as a potential anti-adipogenic lead compound.

## Conclusion

As concluding remarks, the results obtained demonstrated that cuminaldehyde and cuminaldehyde and cuminaldehyde-rich essential oil extracted from *C. comosum* essential oil exhibited both suppression in pre-adipocytes maturation and an increase in lipid turnover; indicative of a potential anti-adipogenic activity. This study can be considered, up to our best of knowledge, as the first report for the anti-adipogenic activities of *C. comosum* essential oil extract and cuminaldehyde. The extract from *C. comosum* and/ or its major component cuminaldehyde can be further investigated for the use as a potential therapeutic agent for the prevention and treatment of obesity.

## Abbreviations

CEBPa: CAAT Enhancer Binding Protein alpha,
PPARg: Peroximal Proliferator-activated receptor gamma.

## Acknowledgements

This research was fulfilled with the competitive grant (1801050130-P) and targeted grant (1801050232-P) from University of Sharjah.

## Author contributions

MGM, AE and SS designed and developed the study. MGM and SS conceived the study and wrote the manuscript. SS was responsible for oil extraction, identification and analysis of fractions and compound. AE, AW, DP, RA and MM were responsible for laboratory work including cell culture, treatment, cell viability assay, Nile Red staining, glucose, cholesterol and triglyceride quantification. AE, AW and DS carried out the RNA extraction and Gene expression experiments. MGM was responsible for data analysis and statistical calculations. MGM, SS interpreted the results and wrote the manuscript. All authors read and approved the final manuscript.

## References

1. WHO (2020) Obesity and overweight.

2. Akil L, Ahmad HA (2011) Relationships between obesity and cardiovascular diseases in four southern states and Colorado. Journal of health care for the poor and underserved 22: 61–72.

3. Viester L, Verhagen EALM, Oude Hengel KM, Koppes LLJ, van der Beek AJ, et al. (2013) The relation between body mass index and musculoskeletal symptoms in the working population. BMC musculoskeletal disorders 14: 238–238.

4. Garg SK, Maurer H, Reed K, Selagamsetty R (2014) Diabetes and cancer: two diseases with obesity as a common risk factor. Diabetes, obesity & metabolism 16: 97–110.

5. Hill JO, Wyatt HR, Peters JC (2012) Energy balance and obesity. Circulation 126: 126–132.

6. Karam J, McFarlane S (2010) Tackling obesity: new therapeutic agents for assisted weight loss. Diabetes, metabolic syndrome and obesity : targets and therapy 3: 95–112.

7. Muir LA, Neeley CK, Meyer KA, Baker NA, Brosius AM, et al. (2016) Adipose tissue fibrosis, hypertrophy, and hyperplasia: Correlations with diabetes in human obesity. Obesity (Silver Spring, Md) 24: 597–605.

8. Park U-H, Jeong H-S, Jo E-Y, Park T, Yoon SK, et al. (2012) Piperine, a Component of Black Pepper, Inhibits Adipogenesis by Antagonizing PPARγ Activity in 3T3-L1 Cells. Journal of Agricultural and Food Chemistry 60: 3853–3860.

9. Wang ZQ, Zhang XH, Yu Y, Tipton RC, Raskin I, et al. (2013) Artemisia scoparia extract attenuates non-alcoholic fatty liver disease in diet-induced obesity mice by enhancing hepatic insulin and AMPK signaling independently of FGF21 pathway. Metabolism: clinical and experimental 62: 1239–1249.

10. Tang Y, Tsao R (2017) Phytochemicals in quinoa and amaranth grains and their antioxidant, anti-inflammatory, and potential health beneficial effects: A review. 1600767 p.

11. Wang Q, Wang S-t, Yang X, You P-p, Zhang W (2015) Myricetin suppresses differentiation of 3 T3-L1 preadipocytes and enhances lipolysis in adipocytes. Nutrition Research 35: 317–327.

12. Zhang T, Sawada K, Yamamoto N, Ashida H (2013) 4-Hydroxyderricin and xanthoangelol from Ashitaba (Angelica keiskei) suppress differentiation of preadiopocytes to adipocytes via AMPK and MAPK pathways. Molecular Nutrition & Food Research 57: 1729–1740.

13. Centre for Mediterranean Cooperation IUCNNRU (2005) A guide to medicinal plants in North Africa: IUCN Centre for Mediterranean Cooperation.

14. Mandaville JP (2011) Bedouin Ethnobotany: Plant Concepts and Uses in a Desert Pastoral World: University of Arizona Press.

15. Badria Farid A, Ameen M, Akl Mohamed R (2007) Evaluation of cytotoxic compounds from Calligonum comosum L. growing in Egypt. Zeitschrift für Naturforschung C. pp. 656.

16. Liu XM, Zakaria MNM, Islam MW, Radhakrishnan R, Ismail A, et al. (2001) Anti-inflammatory and anti-ulcer activity of Calligonum comosum in rats. Fitoterapia 72: 487–491.

17. Iacobellis NS, Lo Cantore P, Capasso F, Senatore F (2005) Antibacterial activity of Cuminum cyminum L. and Carum carvi L. essential oils. J Agric Food Chem 53: 57–61.

18. Soliman SSM, Abouleish M, Abou-Hashem MMM, Hamoda AM, El-Keblawy AA (2019) Lipophilic Metabolites and Anatomical Acclimatization of Cleome amblyocarpa in the Drought and Extra-Water Areas of the Arid Desert of UAE. Plants 8: 132.

19. Soliman S, Mohammad MG, El-Keblawy AA, Omar H, Abouleish M, et al. (2018) Mechanical and phytochemical protection mechanisms of Calligonum comosum in arid deserts. PLOS ONE 13: e0192576.

20. Soliman S, Abouleish MY, Khoder G, Khalid B, Husam H, et al. (2020) The scientific basis of the antibacterial traditional use of Calligonum comosum in UAE. Journal of Herbal Medicine: 100361.

21. Soliman S, Alsaadi A, Youssef E, Khitrov G, Noreddin A, et al. (2017) Calli Essential Oils Synergize with Lawsone against Multidrug Resistant Pathogens. Molecules 22: 2223.

22. Kokoska L, Havlik J, Valterova I, Sovova H, Sajfrtova M, et al. (2008) Comparison of chemical composition and antibacterial activity of Nigella sativa seed essential oils obtained by different extraction methods. J Food Prot 71: 2475–2480.

23. Vishwanath D, Srinivasan H, Patil MS, Seetarama S, Agrawal SK, et al. (2013) Novel method to differentiate 3T3 L1 cells in vitro to produce highly sensitive adipocytes for a GLUT4 mediated glucose uptake using fluorescent glucose analog. Journal of cell communication and signaling 7: 129–140.

24. Ramakrishnan L, Reddy KS, Jailkhani BL (2001) Measurement of Cholesterol and Triglycerides in Dried Serum and the Effect of Storage. Clinical Chemistry 47: 1113–1115.

25. Bakker AJ, Mücke M (2007) Gammopathy interference in clinical chemistry assays: mechanisms, detection and prevention. 45: 1240.

26. Estes BT, Guilak F (2011) Three-dimensional culture systems to induce chondrogenesis of adipose-derived stem cells. Methods in molecular biology (Clifton, NJ) 702: 201–217.

27. El-Serafi AT, Sandeep D, Abdallah S, Lozansson Y, Hamad M, et al. (2019) Paradoxical effects of the epigenetic modifiers 5-aza-deoxycytidine and suberoylanilide hydroxamic acid on adipogenesis. Differentiation 106: 1–8.

28. Tang Q-Q, Zhang J-W, Daniel Lane M (2004) Sequential gene promoter interactions of C/EBPβ, C/EBPα, and PPARγ during adipogenesis. Biochemical and Biophysical Research Communications 319: 235–239.

29. Bergman RN, Stefanovski D, Buchanan TA, Sumner AE, Reynolds JC, et al. (2011) A better index of body adiposity. Obesity (Silver Spring, Md) 19: 1083–1089.

30. Otto TC, Lane MD (2005) Adipose Development: From Stem Cell to Adipocyte. Critical Reviews in Biochemistry and Molecular Biology 40: 229–242.

31. den Hartigh LJ (2019) Conjugated Linoleic Acid Effects on Cancer, Obesity, and Atherosclerosis: A Review of Pre-Clinical and Human Trials with Current Perspectives. Nutrients 11: 370.

32. Camp HS, Ren D, Leff T (2002) Adipogenesis and fat-cell function in obesity and diabetes. Trends in Molecular Medicine 8: 442–447.

33. Jang M-K, Yun Y-R, Kim J-H, Park M-H, Jung MH (2017) Gomisin N inhibits adipogenesis and prevents high-fat diet-induced obesity. Scientific Reports 7: 40345.

34. Harris RBS, Kasser TR, Martin RJ (1986) Dynamics of Recovery of Body Composition After Overfeeding, Food Restriction or Starvation of Mature Female Rats. The Journal of Nutrition 116: 2536–2546.

35. Li KK, Liu CL, Shiu HT, Wong HL, Siu WS, et al. (2016) Cocoa tea (Camellia ptilophylla) water extract inhibits adipocyte differentiation in mouse 3T3-L1 preadipocytes. Scientific reports 6: 20172–20172.

